# Multi-Center Study of Resectable Lung Lesions by Ultra-Deep Sequencing of Targeted Genes in Plasma Cell-Free DNA to Assess Nodule Malignancy and Detect Lung Cancers

**DOI:** 10.1101/453803

**Authors:** Muyun Peng, Yuancai Xie, Xiaohua Li, Youhui Qian, Xiaonian Tu, Xumei Yao, Fangsheng Cheng, Feiyue Xu, Deju Kong, Bing He, Chaoyu Liu, Fengjun Cao, Haoxian Yang, Jiankui He, Fenglei Yu, Chuanbo Xu, Geng Tian

**Author notes:** Corresponding authors: C. Xu,; F. Yu,; G. Tian. Authors contributed equally (Muyun Peng, Yuancai Xie, Xiaohua Li).

## Abstract

**BACKGROUND:** Early detection of lung cancer to allow curative treatment remains challenging. Cell-free circulating tumor DNA (ctDNA) analysis may aid in malignancy assessment and early cancer diagnosis of lung nodules found in screening imagery.

**METHODS:** The multi-center clinical study enrolled 192 patients with operable occupying lung diseases. Plasma ctDNA, white blood cell genomic DNA (gDNA) and tumor tissue gDNA of each patient were analyzed by ultra-deep sequencing to an average of 35,000X of the coding regions of 65 lung cancer-related genes.

**RESULTS:** The cohort consists of a quarter of benign lung diseases and three quarters of cancer patients with all histopathology subtypes. 64% of the cancer patients is at Stage I. Gene mutations detection in tissue gDNA and plasma ctDNA results in a sensitivity of 91% and specificity of 88%. When ctDNA assay was used as the test, the sensitivity was 69% and specificity 96%. As for the lung cancer patients, the assay detected 63%, 83%, 94% and 100%, for Stage I, II, III and IV, respectively. In a linear discriminant analysis, combination of ctDNA, patient age and a panel of serum biomarkers boosted the overall sensitivity to 80% at a specificity of 99%. 29 out of the 65 genes harbored mutations in the lung cancer patients with the largest number found in TP53 (30% plasma and 62% tumor tissue samples) and EGFR (20% and 40%, respectively).

**CONCLUSION:** Plasma ctDNA was analyzed in lung nodule assessment and early cancer detection while an algorithm combining clinical information enhanced the test performance.

## Introduction

Lung cancer is the leading cause of cancer-related deaths, accounting for an estimated 1.6 million deaths each year globally (1). The prognosis of lung cancer is dependent on the stage of diagnosis, with 5-year overall survival rate decreasing dramatically from stage IA (85%) to stage IV disease (6%) (2, 3). This makes it clearly a winning strategy to screen and diagnose lung cancer earlier to saving lives (3).

Current method for lung cancer screening is low-dose computed tomography (LDCT) (4). The National Lung Screening Trial (NLST) demonstrated a 20% reduction in lung cancer mortality for LDCT compared with X ray and a 6.7% all-cause mortality reduction (5). However, the imagery technique often results in indeterminate nodules (6) and the false positive results lead to unnecessary invasive diagnostic procedures and increased deaths from avoidable surgeries (7).

Traditionally, biopsy is used to determine malignancy of lung nodules. This approach has significant limitations as being difficult or even impossible. Molecular tests using pervasive biofluid samples, so called liquid biopsy, are promising and urgently needed in the thoracic clinic, as demonstrated in thyroid nodules assessment where a number of molecular tests are available (8). In pulmonary nodule malignancy assessment, there is also the report of blood proteomics biomarkers by the PANOPTIC team (9).

In theory, ctDNA is exquisitely specific for an individual’s tumor as by definition somatic mutations are identified by their presence in tumor DNA and absence in matched normal DNA. This bypasses the issues related to the false-positivity encountered with other biomarkers, such as protein biomarkers. This specific and promising early detection method has garnered tremendous attention for cancer in general and lung in particular (10,11). The growing interest has drawn attention from international societies such as IASLC that has issued statement regarding liquid biopsy in the management of non-small cell lung cancer (NSCLC) (12).

Early detection of cancer by blood test was shown possible even with microscopic tumor proceeding radiography (13). Most recently, a preliminary analysis of early to mid-stage (stage I-III) lung cancer patients as part of the Circulating Cancer Genome Atlas (CCGA) pan-cancer study was reported (14).

In addition to mutation analysis, ctDNA epigenetics has also been evaluated and researched for early cancer detection. The studies examined DNA methylation regulation (15) or the hypermethylation of the promoter regions of genes (16) for the potential biomarkers of lung cancer detection.

In this study, we report a multi-center clinical trial result on ultra-deep sequencing in patients that undergo surgical resection either benign nodules or with different stages of lung cancer.

## Patients & Methods

### Clinical Centers and Patients

The clinical trial (ClinicalTrial.gov NCT03081741) at four Tier A hospitals in China recruited patients with lung occupying diseases to be treated by surgery. Patients with benign lung nodules or cancers of Stage I to III planned for surgical resection were eligible for inclusion (Table S1). After informed consenting, biological samples (two tubes of 10mL peripheral blood prior to surgery collected in a cell-free DNA (cfDNA) BCT blood collection tube (Streck, Omaha, US) and 10 slides of formalin fixed paraffin embedded (FFPE) tissues from surgery), and related clinical data including serum biomarkers were collected.

### Sequencing Analysis

The above collected DNA samples were analyzed by our proprietary Sec-Seq technique as described previously (17). Briefly, blood sample were processed to separate the plasma from blood cells by centrifugation. cfDNA, gDNA of WBC and FFPE were extracted using QIAamp Circulating Nucleic Aid Kit, DNA mini kit, DNA FFPE Tissue kit, respectively (Qiagen, Hilden, Germany). The concentration of extracted DNA was measured using Qubit 3.0 dsDNA high-sensitivity assay (Life Technologies, Carlsbad, CA).

Capture probes were designed for 65 cancer-associated genes covering 241kb genomic regions (Table S2) and synthesized by IDT (MI, USA). Indexed libraries were constructed using KAPA HyperPlus Kit (KK8514). Barcoding was employed to reduce noise. Post-capture multiplexed libraries were amplified with Illumina backbone primers for 16 cycles of PCR using 1× KAPA HiFi Hot Start Ready Mix and sequenced on Illumina NovaSeq platform (Illumina, CA, USA) at Novogen (Nanjing, China).

Horizon’s Partners Spike-in control (Horizon, Cambridge, UK) was used in a serial dilution (from 0.0005 to 1) using the wild-type reference genome and the provided reference standard. Reference variants include EGFR (L858R, T790M), KRAS (G12D), NRAS (Q61K, A59T) and PIK3CA (E545K), or NA12878 (12 mutations) and NA24385 (29 mutations).

### Bioinformatics Data Analysis

Reads with quality score <30 or havinh > 5% of positions differ from the rest of the reads targeting the same region were removed. The results were then mapped to the human reference genome (hg19) using BWA (v0.7.15-r1140). We used the start mapping positions, the length and the dual barcode on both side of merged paired-end fragments to form reads groups amplified from every primary cfDNA molecules and to identify incorrect base produced due to PCR errors.

Variant calling for single nucleotide variation (SNV) or insertion/deletion (INDEL) was performed using samtools mpileup tool (v1.3.1). For ctDNA samples, a variant was selected as a candidate somatic mutation when (1) two distinct paired reads (each redundantly sequenced at least three times) contained the mutation, (2) effective reads depth >500 (captured primary cfDNA molecules > 500) and (3) the corresponding allele frequency in WBC is less than 1%.

### Mutation Annotation and Classification

The variants were called by SnpEff (v4.3o), and annotated by COSMIC (v85), ExAc, ClinVar, and 1000 Genome. The following variants were eliminated: (1) intergenomic or intronic (except for splicing junction); (2) synonymous; (3) variant allele frequency (VAF) <0.2% in ctDNA or <1% in FFPE samples. Previously reported and confirmed pathogenic mutations in the clinical samples of lung cancer of all human races and ethnicities will be considered as lung cancer related.

### Statistical Methods

Data were summarized using descriptive statistics. Fisher’s exact test was used to compare any two subgroups. Wilcoxon rank sum test was used to compare median age between any two subgroups (Stage I, II, III) or mutant groups (mutant vs wild type).

Linear discriminant analysis (LDA) was performed on improving mutation analysis. The model considers the age of the patients, ctDNA mutations, and the serum biomarkers. It was developed using the 10 fold cross validation by dividing the samples into training and validation subsets. The test sensitivity and specificity was calculated, and AUROC was plotted.

## Results

### Patient Demographic and Clinical Analysis

In total, 192 patients with pulmonary space occupying lesions (136 malignant and 56 benign) pathologically diagnosed and surgically treated were included in this analysis. These patients were recruited from 4 clinical sites: Xiangya No.2 Hospital in Changsha, Hunan Province, Beijing University Shenzhen Hospital, Shenzhen, Huizhou People’s Hospital, Huizhou, and No.2 People’s Hospital in Shenzhen, Guangdong Province. Though excluded but due to delayed pathological determination, three Stage IV patients were enrolled and hence analyzed.

The average age is 56.5 (range 26-79) years and male proportion is 59%. These numbers are 50.1 (26-73) years and 55% for the benign group, 59.1 (27-79) years and 60% for lung cancer group with a statistically significant difference. No difference was found between the two groups for smoke status (32% vs. 35%) and family history (7% vs. 8%) (Table 1).

**Table 1.** Patient Demographics.

|  | Non-Cancer | Cancer | Total |
| --- | --- | --- | --- |
| Patients | n=56 | n=136 | n=192 |
| Sex |  |  |  |
| Male | 31 (55.4%) | 82 (60.3%) | 113 (58.9%) |
| Female | 25 (44.6%) | 54 (39.7%) | 79 (41.1%) |
| Average Age (Range) | 50.1 (26-73) | 59.1 (27-79) | 56.5 (26-79) |
| Smoking History |  |  |  |
| Yes | 18 (32.1%) | 48 (35.3%) | 66 (34.4%) |
| No | 38 (67.9%) | 88 (64.7%) | 126 (65.6%) |
| Family History |  |  |  |
| Yes | 4 (7.1%) | 11 (8.1%) | 15 (7.8%) |
| No | 52 (92.9%) | 125 (91.9%) | 177 (92.2%) |
| Lung Nodule Size (mm, Avg. Max Dimension) | 23.3 | 29.4 | 27.6 |

For the benign lesions, the most common diseases diagnosed are pneumonia (n=14, 25%), tuberculosis (n=12, 21%), pulmonary fibrosis (n=4, 7%), and necrosis granuloma (n=3, 5%).

Lung cancer distribution was 87, 29, and 17 in Stages I, II and III, respectively. The average size of the nodules for lung cancer patients is 2.9 (range 0.5 – 9.0) cm, and for each subgroup: (1) Stage I: 2.2 (0.5 - 4.0) cm; (2) Stage II: 3.8 (1.0 - 7.0) cm; and (3) Stage III: 5.0 (1.3 - 9.0) cm (Table 2). The average size of the nodules in the benign group is 2.3 (0.3 - 6.0) cm which is statistically smaller than that of the malignant group.

**Table 2.** Lung Cancer Histological Diagnosis and Stages.

|  | Number of Pts (percentage) | Average Nodule<br>Size (mm, largest<br>dimension) |
| --- | --- | --- |
| Pathological Diagnosis |  |  |
| Adenocarcinoma | 100 (73.5%) | 24.3 |
| Squameous Carcinoma | 28 (20.6%) | 49.8 |
| Small Cell Lung Cancer | 1 (0.7%) | 18 |
| Others | 7 (5.2%) | 22.3 |
| Clinical Stage |  |  |
| I | 87 (64.0%) | 22 |
| II | 29 (21.3%) | 39 |
| III | 17 (12.5%) | 52.1 |
| IV | 3 (2.2%) | 21 |

All of the patients are with solitary pulmonary nodules, except for two patients who had two nodules each that are malignant.

### Genetic Profiling and Mutation Burden

For each patient, three biospecimen samples including plasma ctDNA, WBC and FFPE tumor tissue were sequenced. For ctDNA, the average sequencing depth is 35000 with 1350 unique reads after deduplication.

In total, 312 occurrences and 274 unique somatic mutations were found in 29 genes from either plasma ctDNA or tissue DNA in 120 cancer and 5 benign cases. No mutations were found in 14 lung cancers in either blood or tissue samples.

In the benign lesions, 2 out of 56 patients (3.6%) had 4 non-driver gene mutations in ctDNA, and 3 patients (5.4%) had 2 non-driver mutations in FFPE samples.

Among the lung cancer patients, 88% (120 out of 136 patients) were found to harbor at least one mutation in ctDNA or tumor tissue. When analyzed by stage of cancer, the class, that is whether driver or non-driver mutation, and number of mutations increase as the stage advances (Figure 1).

**Figure 1.**
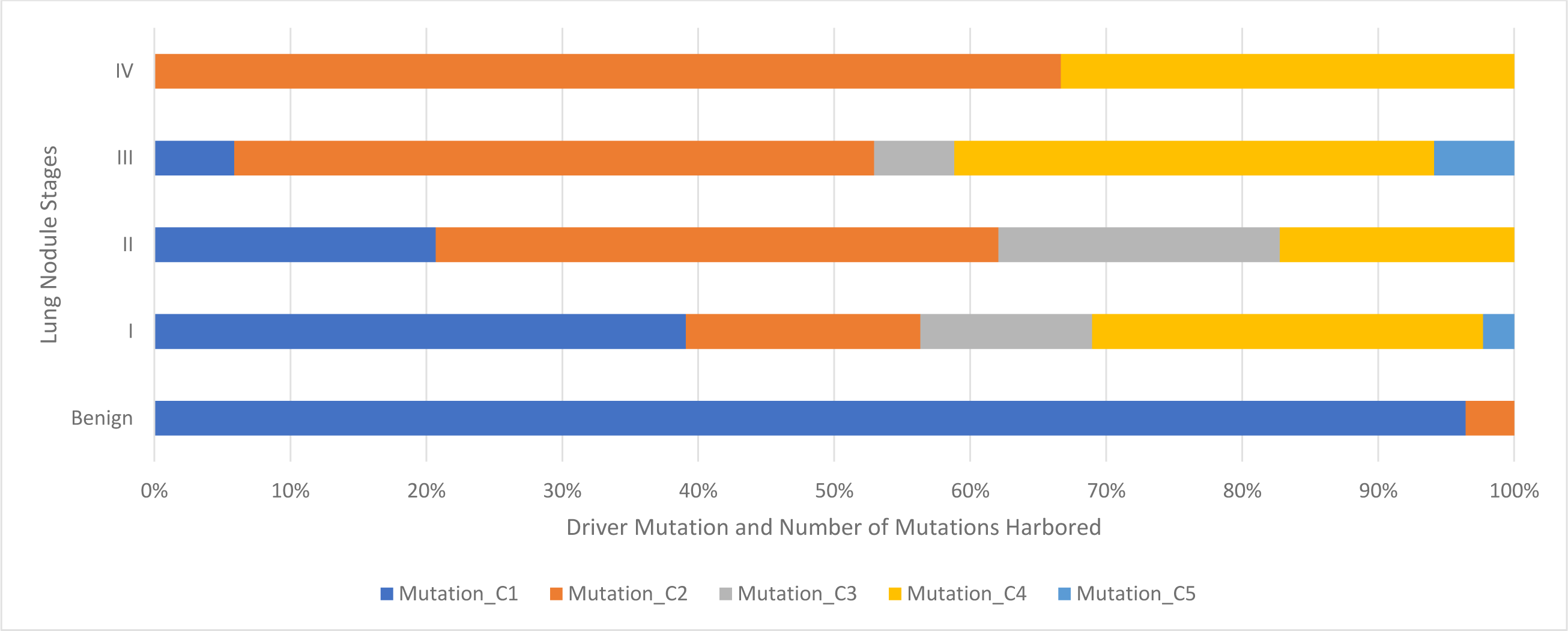
Lung Cancer Gene Mutation Burden in Benign and Various Stages of Patients.

Notes: Mutation types: **C1**. No mutation found; **C2**. 1 non-driver mutation found; **C3**. 1 driver mutation found; **C4**. Both driver mutation and non-driver mutation found; **C5**. More than 1 non-driver mutation found.

Mutations were found in 9 known lung cancer genes of ALK, BRAF, EGFR, HER2, KRAS, MET, NRAS, PIK3CA, and ROS1 (Fig. 2) out of the 12 genes defined as drivers for lung cancer (Ref. 18). The most commonly mutated genes in the lung cancer patients are TP53 (44%) and EGFR (35%) (Figure S1).

**Figure 2.**
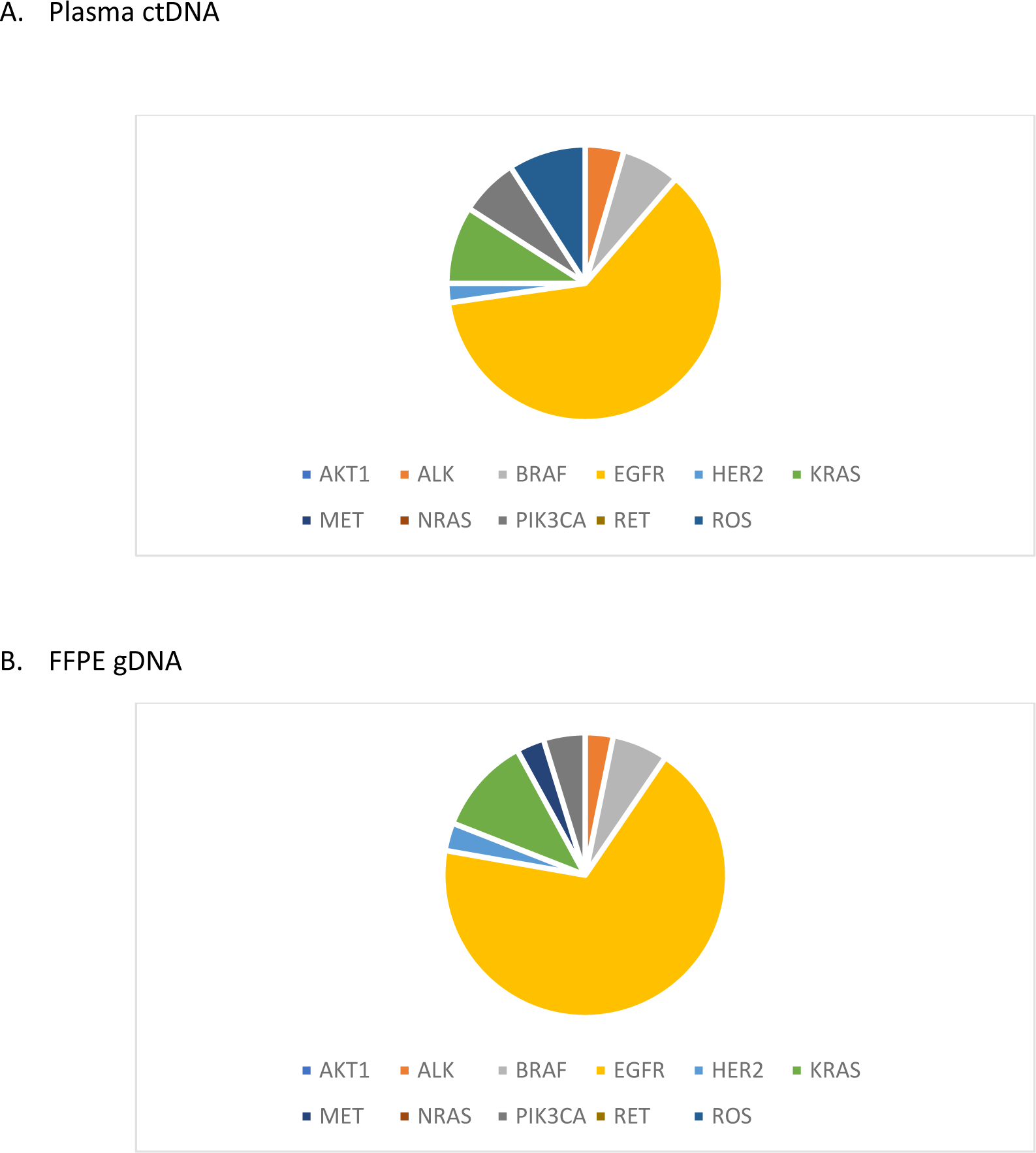
Driver Mutation Distribution in Lung Cancer Patients.

The most commonly occurred mutation is EGFR L858R (found in 24 samples) followed by EGFR exon 19 deletion (in 15 samples). The largest number of mutations was found in a patient who harbored 15 mutations.

### Concordance between ctDNA and Tumor Tissue

Concordance is defined as at least one gene mutation is the same in both the plasma ctDNA and the FFPE tumor tissue gDNA of a patient. When completely no mutations were found in both the blood ctDNA and tissue gDNA, it is also considered as concordant of the two samples.

Among the 136 malignant cases, the overall concordance rate is 37%. Among the concordant patients, 35 of them share at least one mutation in blood and tissue samples, and 15 of them had no mutations in either the ctDNA or tissue samples. Concordance was higher in the driver genes, 46%. The shared mutation rate increases as the stage of the cancer advances: 32%, 78%, 50%, and 67% at Stage I, II, III, and IV, respectively (Figure S2).

### Comparison with Serum Biomarkers

A panel of six serum protein tumor biomarkers was also analyzed which has a sensitivity of 51% with a specificity of 83% (Table 3). These markers include NSE, CYFRA 21-1, CEA, ProGRP, CA-125, and SCC. When the most sensitive marker of CYFRA 21-1 was considered alone, the sensitivity was merely 25% at a specificity of 95% (Table 3).

**Table 3.**
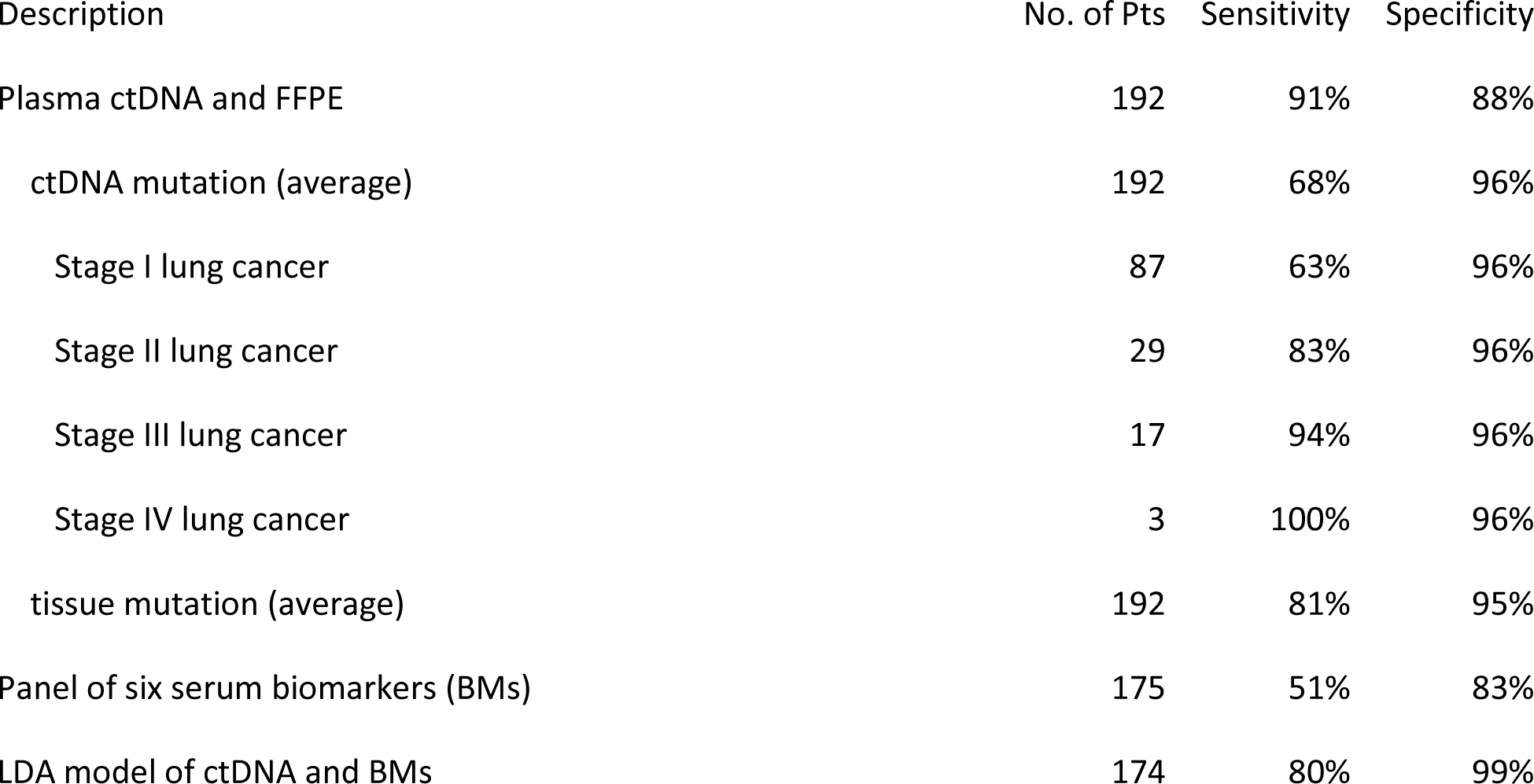
Lung nodule malignancy prediction diagnostic test Performance.

| Description | No. of Pts | Sensitivity | Specificity |
| --- | --- | --- | --- |
| Plasma ctDNA and FFPE | 192 | 91% | 88% |
| ctDNA mutation (average) | 192 | 68% | 96% |
| Stage I lung cancer | 87 | 63% | 96% |
| Stage II lung cancer | 29 | 83% | 96% |
| Stage III lung cancer | 17 | 94% | 96% |
| Stage IV lung cancer | 3 | 100% | 96% |
| tissue mutation (average) | 192 | 81% | 95% |
| Panel of six serum biomarkers (BMs) | 175 | 51% | 83% |
| LDA model of ctDNA and BMs | 174 | 80% | 99% |

In comparison, the profiling by ctDNA showed a higher sensitivity in detecting lung cancer. The sensitivity increases as the stage advances and ctDNA outperform the serum biomarkers in all stages (Figure 3).

**Figure 3.**
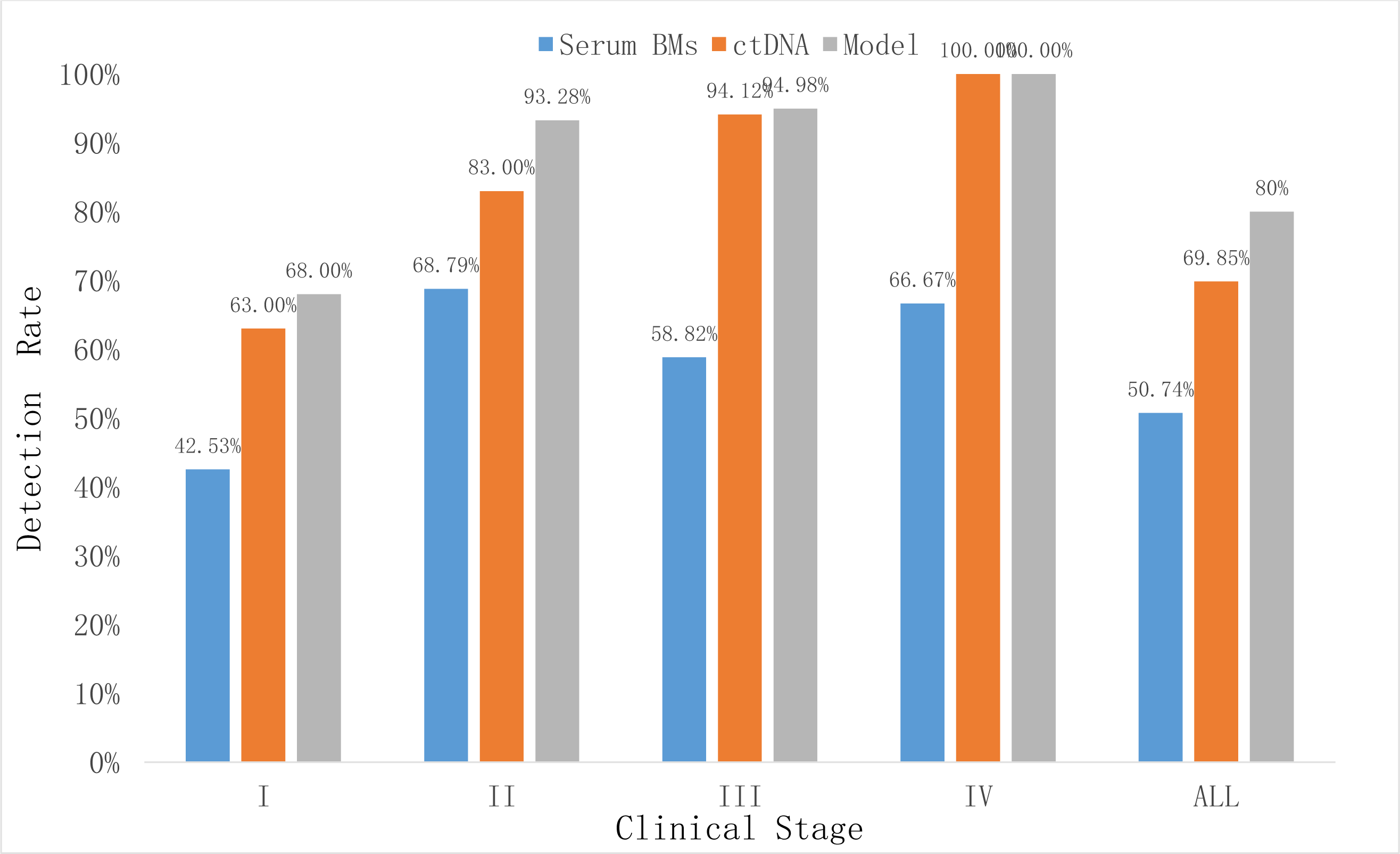
Lung Cancer Detection Rate by ctDNA, Serum Biomarkers and LDA Model per Stages.

Note: Model = LDA (Linear Discriminant Analysis) model.

### Malignancy Assessment and Lung Cancer Tests

The mutation profiling classifies the lung nodules from benign to malignant (Figure 1). The benign samples harbor very little to no mutations. The cancer patients have increasing level of mutations both in terms of category and numbers as stage progresses. From Stage I to IV, there are more mutations found and they are more often in driver genes.

For lung cancer detection, the overall sensitivity of plasma ctDNA was 68% at the specificity of 96%. Slightly higher detection rate was observed in tissue DNA which reached 78% (Table 3). According to cancer stages, the sensitivity rate is 60%, 69%, 94%, and 100% for Stage I, II, III, and IV, respectively (Fig. 3).

We further conducted a linear discriminant analysis where patient age, smoking status, and serum protein markers were considered (Figure S3). The combined model of ctDNA mutations and serum biomarkers improved the sensitivity and specificity to 80% and 99% respectively (Table 3).

## Discussion

Lung cancer screening recommended by the guidelines targets populations at high risk of developing lung cancer, such as patients aged above 55 years, heavy smokers and having chronic obstructive pulmonary disease (COPD) and with family history of lung cancer. In general, by definition of 50 – 500 mm^3^ (19), screening by imagery techniques could result in about 30-40% indeterminate nodules that need to be further evaluated. In our cohort, we have >60% nodules in this size range making it an urgent need for malignancy assessment.

Our patient population drawn from major hospitals in southern and central China has a median age of 56.5 years, just above the threshold for lung cancer screening. The clinical outcome confirmed the age as a risk factor for lung cancer: the median age of the patients with benign nodules (50.1 years) was about ten years younger than those having lung cancer (median age of 59.1 years).

Smoking status and family history were not significantly different between the benign and the malignant cases in our cohort. One possible explanation could be that the impact of smoking should be considered more broadly than tabaco smoking alone. For example, the time spent on Chinese style cooking could be a potential risk. Unfortunately, no such data were collected. As a proxy, however, the gender distribution difference could be explored since female usually spends more time in the kitchen. While we have more male patients in the cancer group (about 60%), the opposite is true for the benign group which has less (about 55%).

It is reported that tumor burden of lung cancer corresponds to its size (20). Our data confirms that the size of the nodule relates to malignancy and progression. The average tumor size increases from 22mm in Stage I to 38mm in Stage II and to 50mm in Stage III (Table 2). However, there is no significant difference between a benign nodule and that of the Stage I cancer (Table 1 and 2).

The concordance rate between tissue and corresponding plasma ctDNA also reflects the challenge of early cancer liquid biopsy. Our study is heavy in Stage I patients (64%, Table 2) which has a rate of 32% and causes the overall rate of 36%. The rate increases as stage advances – up to the highest of 78% and the average of Stage I-III is 53%. In CCGA, a set of 73 early to mid-stage (stage I-III) lung cancer samples showed a similar rate of 59% (CI = 47-70%) (13). In another very small study of 31 paired lung cancer tissues and plasma DNA samples with 10,000-fold ctDNA sequencing depth, the concordance of mutation between tumor tissue DNA and ctDNA was merely 3.9% (21). Ours is more like CCGA in that we both sequenced ctDNA to the depth >40,000X. Another meta-analysis has also put the pooled sensitivity in the range of 60%’s (22).

The gene mutations shared between the plasma ctDNA and the FFPE tumor tissue increase as the lung cancer stage advances. This is in alignment with the previous report that Stage IV tumor has the highest concordance (23), as well as that as the tumor is getting larger, the amount of DNA fragments it sheds into the blood stream will increase (20,24).

Although somatic mutations showed strong feasibility of detecting malignancy and staging the cancer by plasma ctDNA, there more factors to be considered. Therefore, combining clinical and genomic features improved the test performance as shown by our LDA modeling (Table 3). Integrated classifiers have also been explored in terms of plasma protein biomarkers such as the ones tested by some panel components in our cohort (25).

Liquid biopsy starts moving into cancer clinics in therapy selection (26, reviewed in 27). For early detection of lung cancer, one study involving 60 NSCLC patients, 40 COPD patients and 40 healthy controls showed that serum cfDNA concentrations and integrity may be valuable (27). Another study identified 17 miRNA species in the exosomes of the blood that are differentially expressed in cancer (both NSCLC and SCLC) and controls (27). The potential use of ctDNA for early detection of other cancers has also been reported (29,30).

There are, however, a number of limitation and challenges. First of all, the sample sizes of the early detection studies are usually small especially the number of healthy controls. Second, ctDNA amount correlates with cancer stage (31). Therefore, the consensus is that ultra-deep sequencing of 40,000X is required to detect the low frequency mutations in the 10mL blood. Finally, the less than expected driver mutation concordance between ctDNA and tumor DNA may reflect genetic heterogeneity and indicate tumor evolution (32) suggesting that other types of genes and mutations should be considered as well.

Some biomarkers for lung cancer in blood can be present in patients without identifiable radiological nodules and thus precede the apparition of these nodules. We are continuing with long term follow-up to study these biomarkers. We also continue working on technical improvement of the assay as well as the clinical validation.

## Statements

### Contributorship

MP, YX, YQ, FC, HY, FY, GT participated in patient recruitment; FC, CL, CX participated in clinical trial and patient sample management; XL, XY, FX participated in genomic sequencing; XT, DK, BH, JH, CX participated in bioinformatics data analysis; and CX, XL, FY, GT participated in manuscript writing and review.

## Acknowledgement

The authors would like to thank the nurses in the Department of Thoracic Surgery, No.2 Xiangya Hospital, Thoracic Department, Peking University Shenzhen Hospital, Department of Oncology, Shenzhen Second People’s Hospital, Oncology Center and Department of Thoracic Surgery, Sun Yat-sen University Cancer Center for their work in enrolling the patients, collecting biospecimen samples, and related clinical data. We would like to thank the laboratory staff at Vienomics for processing the samples and conducting the sequencing procedures. Finally but not the least, we would like to express our highest appreciation of the participation and contribution of our patients in the study.

